# MSBooster: Improving Peptide Identification Rates using Deep Learning-Based Features

**DOI:** 10.1101/2022.10.19.512904

**Authors:** Kevin L Yang, Fengchao Yu, Guo Ci Teo, Vadim Demichev, Markus Ralser, Alexey I Nesvizhskii

## Abstract

Peptide identification in liquid chromatography-tandem mass spectrometry (LC-MS/MS) experiments relies on computational algorithms for matching acquired MS/MS spectra against sequences of candidate peptides using database search tools, such as MSFragger. Here, we present a new tool, MSBooster, for rescoring peptide-to-spectrum matches using additional features incorporating deep learning-based predictions of peptide properties, such as LC retention time, ion mobility, and MS/MS spectra. We demonstrate the utility of MSBooster, in tandem with MSFragger and Percolator, in several different workflows, including nonspecific searches (immunopeptidomics), direct identification of peptides from data independent acquisition data, single-cell proteomics, and data generated on an ion mobility separation-enabled timsTOF MS platform. MSBooster is fast, robust, and fully integrated into the widely used FragPipe computational platform.

## Introduction

Liquid chromatography-tandem mass spectrometry (LC-MS/MS) is an established, widely used high-throughput method for elucidating the proteome [1]. In the typical LC-MS proteomic workflow, proteins are extracted from the samples and digested into peptides, most commonly using trypsin, which cleaves after lysine and arginine residues. For complex samples, if a high depth of protein identification is required, the workflows are combined with fractionation or enrichment techniques (e.g., to increase the detection of phosphorylated peptides). The peptide preparations are then separated using LC coupled online to a mass spectrometer, and the peptides eluding from the LC column are ionized and transferred to the gas phase. The mass-to-charge (m/z) values of all peptide ions from all peptides eluding from the LC column at a particular time (referred to as “retention time”, RT) are measured using the first stage of MS, generating the so-called MS1 spectra. These spectra contain the mass-to-charge (m/z) values of all detectable ions and their intensities. Optionally, ions can also be separated using ion mobility (IM) (available on, e.g., the Bruker’s timsTOF MS platform [2]). In the second stage of MS analysis, selected (typically the most intense) peptide ions are subjected to isolation and fragmentation to break the peptide bond [3] (data independent acquisition (DDA) approach). Alternatively, all peptide ions within a wider (e.g., 10 Da) window of m/z values (data independent acquisition, DIA) [4] or in a continuous quadrupole scan of a particular window size [5] are selected for simultaneous fragmentation. The resulting MS/MS spectra, whether generated in the DIA or DDA mode, contain m/z values, intensities and IM values (when applicable) of all observed fragment ions (e.g., y- and b-ions when using higher energy collisional dissociation, HCD, fragmentation) for the precursor peptides subjected to MS/MS.

The acquired MS/MS spectra, along with their corresponding precursor peptide masses (DDA) or mass windows (DIA), and their RT and IM values, are used to identify the sequences of the peptides that generated the spectra [6]. This is typically done using the sequence database search approach (when using DIA data, optionally with a peptide-fragment deconvolution step to generate pseudo-MS/MS spectra from the original, multiplex MS/MS scans [7]). Computational tools such as MSFragger [8], SEQUEST [9], Andromeda [10], MASCOT [11], MetaMorpheus [12] and Comet [13] compare each experimental MS/MS spectrum against a set of theoretical m/z values of fragments calculated for each candidate peptide based on the provided protein sequence database and assign a score for each peptide-to-spectrum match (PSM). Not every top scoring PSM is a correct identification, due to e.g., noise in the spectra or true peptide sequences missing in the provided protein sequence database [6, 14, 15]. To assist with downstream false discovery rate (FDR) control, decoys (e.g., shuffled or reversed versions of sequences from the “target” protein database) are typically added [6, 16, 17]. The output from the search engines (i.e. the list of PSMs) is used as input to computational post-processing tools such as PeptideProphet [6, 18, 19] and Percolator [20, 21], which combine various search engine scores (e.g., hyperscore, expectation value) and other properties that are useful for discrimination (e.g., mass difference between the theoretical mass of the peptide and that derived from the measured m/z value for that MS/MS scan). The differences in the distributions of scores for decoy peptides versus those of target peptides are used as part of the modeling process to determine the optimal combination of individual features, as well as to calculate posterior probabilities of correct identification and estimate FDR. These tools significantly boost the sensitivity of peptide and protein identification at a fixed FDR compared with filtering the data using individual scores reported by the search engine [6].

Although tools such as PeptideProphet and Percolator are now a part of many computational pipelines, including FragPipe, they do not incorporate prior knowledge regarding peptide separation coordinates (RT, IM) or fragment ion intensities. High confidence PSMs from previously published studies are stored in public repositories and can be leveraged via spectral library searching [22-27], in which known fragment ion intensities help differentiate true from false PSMs. However, relying on experimentally derived spectral libraries is often limiting, as these libraries are inherently incomplete. For instance, protein expression varies from biological condition to condition, cell type to cell type, and genetic background to background, so that libraries can be incomplete even for organisms with large amounts of previous MS/MS data available. Thus, approaches for predicting MS/MS spectra [28] and using predicted spectra from available protein sequence data to improve the sensitivity of peptide identification in LC-MS/MS proteomics have been explored [29-31]. The difference between the experimental and predicted retention times is also known to provide additional discriminating power [32-35] (and it has been incorporated into PeptideProphet modeling [36]). However, the use of RT and MS/MS spectral predictions was initially limited, in part because of the limitations of first-generation prediction algorithms.

More recently, however, a wave of deep learning (DL) models have been trained to predict the physicochemical properties of peptides and MS/MS spectra [37-42]. By training on millions of available peptides, these models can learn general rules to make accurate predictions for new peptides, assuming they are not vastly different from those on which the models were trained. The use of DL-based RT and spectral predictions has been shown to be particularly useful for DIA data analysis [43-45], and for improving the identification rates in immunopeptidome studies concerned with the analysis of human leukocyte antigen (HLA) binding peptides [46-49]. Unfortunately, current PSM rescoring tools that take advantage of DL-based predictions may be difficult for some users to adopt. For example, MaxQuant with Prosit rescoring requires users to upload their database search results to a web server. Rescoring may be performed locally if the users have GPU access, which is not always the case. DeepRescore [50] requires Docker and Nextflow, which may be difficult for users with less computational experience to install.

Here, we present a DL-based PSM rescoring tool MSBooster, a new addition to the widely used FragPipe computational platform, that fully automates the use of DL predictions for improved peptide and protein identification. MSBooster uses DIA-NN [43] (included in FragPipe) to predict the RT, IM and MS/MS spectra of peptides, followed by the generation of additional features for PSM rescoring with Percolator [20]. Importantly, in our approach predictions are performed only for a relatively small set of candidate peptides (e.g., the top 1-5 highest scoring candidates for each MS/MS spectrum). Thus, predictions can be done for each dataset on-the-fly, without the need for a time-consuming full spectral library prediction. We demonstrate the flexibility of MSBooster and its performance in several different workflows, including nonspecific searches (HLA immunopeptidome), DIA quantitative proteomics, single-cell proteomics, and data generated on an IM-enabled timsTOF MS platform. We also explored the behavior of spectral and RT features in the analysis of single cell proteomics data and investigated the potential benefits of using multiple correlated similarity metrics in Percolator. Finally, we assess and discuss the utility of incorporating IM predictions into PSM rescoring.

## Results

### MSBooster and FragPipe computational workflow

We implemented MSBooster as a stand-alone tool that was also fully integrated into FragPipe. FragPipe (https://fragpipe.nesvilab.org/) is a comprehensive computational platform that automates all steps of proteomic analysis, including peptide-spectrum matching with MSFragger [8, 51, 52], PSM validation with PeptideProphet [19] or Percolator [20], protein inference with ProteinProphet [53], and FDR filtering (by default 1% FDR at the PSM, ion, peptide, and protein levels) using Philosopher [54]. FragPipe supports the generation of spectral libraries (using EasyPQP [44]) and the extraction of quantification from DIA data (using DIA-NN [43, 44]). DIA-Umpire [7] is included in FragPipe as one of the modules to generate pseudo-MS/MS spectra from the DIA data. Alternatively, peptides can be identified directly from DIA data using MSFragger-DIA (manuscript in preparation). FragPipe has an easy-to-use graphical user interface (GUI) and includes a data visualization module (FragPipe-PDV), which is an extension of a previously described PDV viewer [55].

Within FragPipe, MSBooster is positioned between MSFragger and Percolator (**Fig 1a**) and is enabled by default in most FragPipe analysis workflows (see **Methods** section for details). In a typical workflow, MSFragger performs the database search, and reports the list of PSMs and associated search scores in a “.pin” file (**Fig 1b**). MSBooster extracts the set of peptides reported in the pin file and creates an input file for a DL-based prediction model (DIA-NN), which in turn generates predictions of the physicochemical properties of peptides, such as RT and IM values, and MS/MS spectra. MSBooster then generates features based on the agreement between the experimental and predicted values and adds them to the original pin files. Finally, it passes these extended pin files to Percolator, which learns a linear support vector machine (SVM) to differentiate true targets from decoys. Percolator assigns a Percolator SVM score and then a posterior error probability to each PSM. DL-based predictions are done for a limited number of peptide candidates: by default, either for a single (top scoring) peptide per MS/MS spectrum (DDA data), top 3 (narrow window GFP-DIA data) or top 5 (conventional DIA data) when using MSFragger-DIA. Thus, in most cases, MSBooster resulted in only a minor increase in the overall computational run time. The timing of the MSBooster steps for the various datasets is shown in **Supplemental Fig 1**.

**Figure 1.**
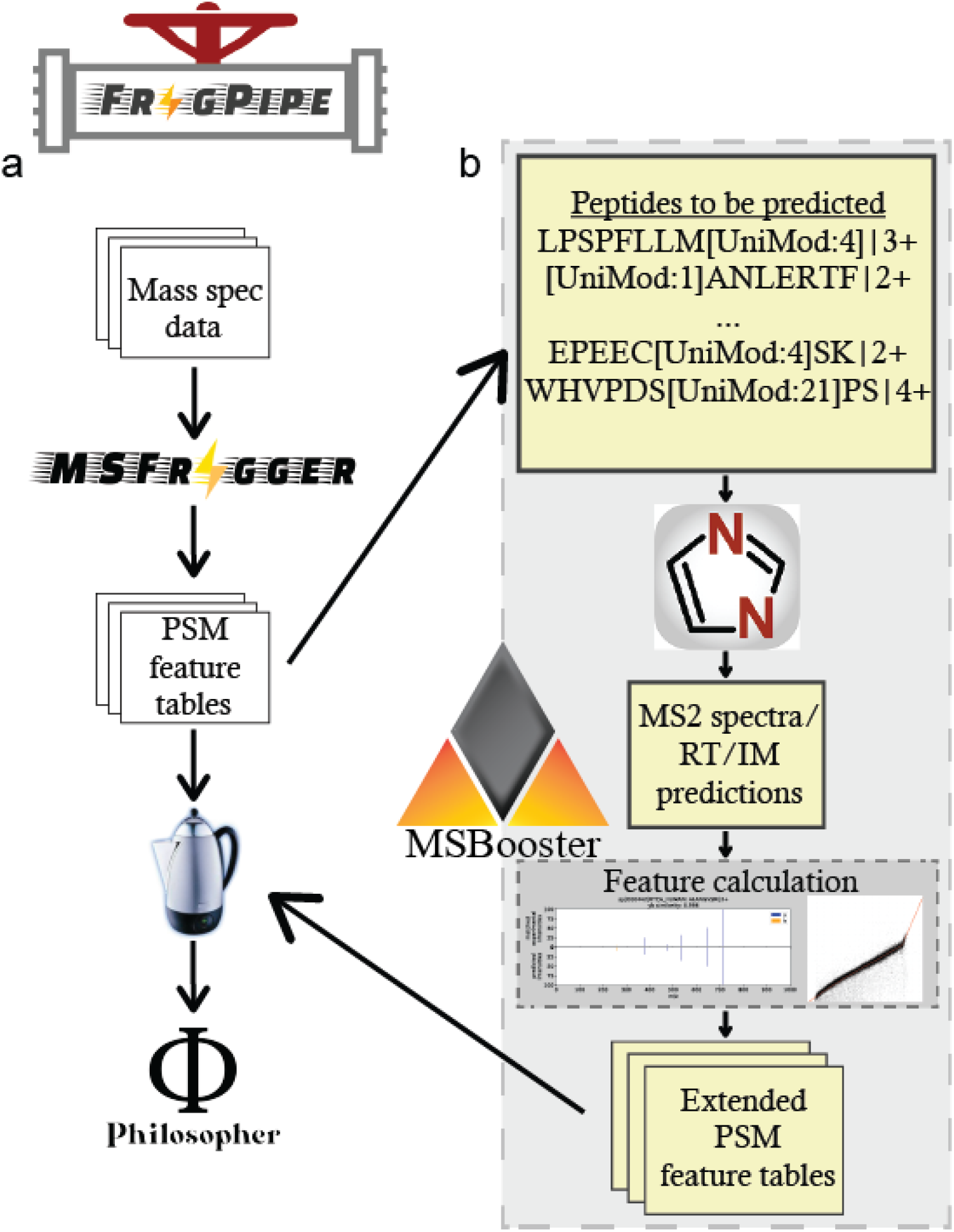
MSBooster workflow. The original workflow without MSBooster (a) and new workflow with MSBooster (b) are depicted. Files generated by MSBooster are depicted in yellow.

### HLA peptide identification

Immunopeptidomics, that is, methods that identify and quantify peptides that are presented as antigens by antigen presenting cells (APCs), are increasingly required in biomedicine but are associated with computational challenges. Human leukocyte antigen (HLA) peptidome data is a promising candidate for DL-based rescoring owing to an expanded nonenzymatic search space, resulting in a higher probability of a high-scoring false match. Because certain major histocompatibility complexes (MHCs) preferentially bind certain peptide motifs, this represents a system in which we know what kinds of peptides should be identified based on their sequences, allowing us to monitor whether MSBooster correctly promotes true target PSMs. To demonstrate the performance of MSBooster with MS/MS spectral and RT-based rescoring on HLA peptides, three fractions of an A*02:01 monoallelic cell line [56] were processed using different combinations of features in MSBooster. Spectral and RT features increased the number of identified peptides by 20.4% and 16.6% respectively, whereas the combination of the two feature types increased the number of identifications by 31.4% (**Fig 2a**). For similar data, Wilhelm et al. showed an average increase of 150% in peptide identification when using MaxQuant coupled with Prosit rescoring [47]. However, this may simply reflect the moderate performance of MaxQuant in nonspecific searches, as noted in [57]. In contrast, even without DL-based rescoring, MSFragger provided multiple and more discriminative scores (**Supplementary Fig 2a**). To illustrate this further, we rescored PSMs using only the hyperscore - one of the main scores from MSFragger - as a starting point (**Supplementary Fig 3a**). Adding spectral and RT features to the hyperscore provided a 183.8% increase in the number of identified peptides. Importantly, adding other features reported by MSFragger (including the mass accuracy and the expectation value) also gives a 161.5% boost compared to using MSFragger’s hyperscore alone. Using all features (all MSFragger computed and all DL-based from MSBooster) results in a 243.7% boost compared with using the hyperscore alone.

**Figure 2.**
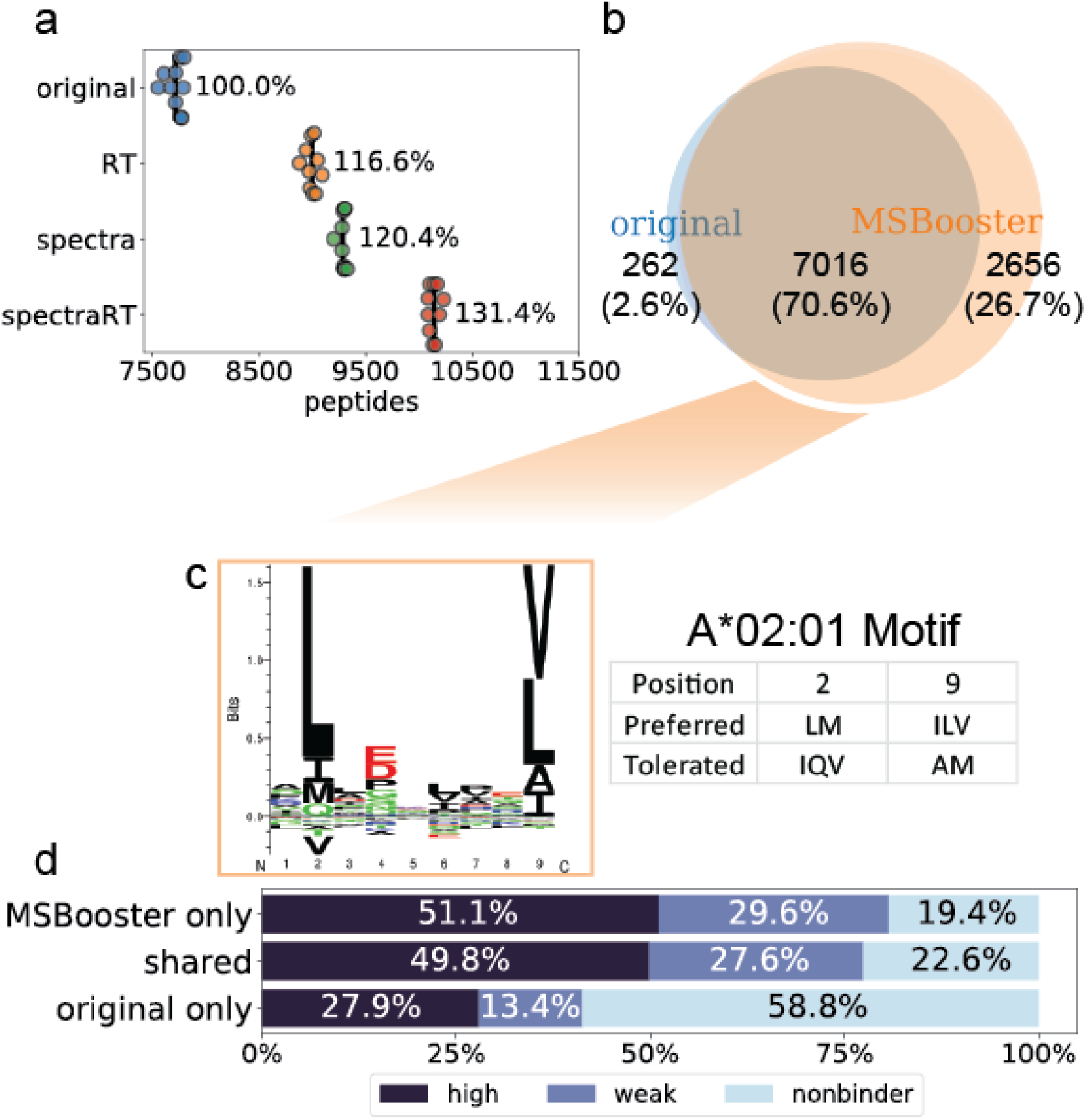
HLA immunopeptidome dataset. (a) Swarmplot of the number of HLA peptides reported at 1% FDR using the MSFragger pin files (original), files with the spectral similarity feature added (spectra), retention time similarity feature (RT), or both types of features (spectraRT). Each dot represents the number reported for each of 10 Percolator runs. Black lines show the average number of peptides reported across 10 Percolator runs. (b) Venn diagram of HLA peptides between lengths 7 and 12 when using either original MSFragger features or with additional deep learning-based features. (c) GibbsCluster-generated motif assigned to the MSBooster-specific peptide subset from (b). The A*02:01 motif was collected from the Immune Epitope Database. (d) Percent of peptides from each subset of (b) that are predicted by NetMHC 4.0 to bind the A*02:01 serotype. Strength of the ligand binding decreases from “high” to “weak” to “nonbinder”.

While most reported peptides passing 1% peptide-level FDR were shared regardless of MSBooster was used (7016 peptides), adding MSBooster resulted in 2656 more identified peptides while only losing 262 (**Fig 2b**). To verify that the added peptides were credible, we identified their HLA sequence motifs using GibbsCluster [58] (**Fig 2c, Supplementary Fig 3b**). MHC binding in the A*02:01 cell line relies on anchors at position 2 and the C-terminus, according to the Immune Epitope Database [59]. The 262 peptides lost after rescoring, when used as input in the motif analysis tool, produced two clusters. The first cluster of 132 peptides followed the expected motif, but the second cluster of 60 peptides was not enriched for the expected amino acids at position 2 (**Supplementary Fig 3b**). Therefore, many of the peptides removed with the help of MSBooster were likely false positives. In contrast, peptides gained with MSBooster generated one cluster of 2533 peptides that faithfully followed the expected sequence motif for the cell line (**Fig 2c**). To further validate the new peptides, we examined their binding affinities with A*02:01 MHC using predictions from NetMHC [60] (**Fig 2d**). We found that 80.7% of the gained peptides and 77.4% of the shared peptides were predicted binders, while this percentage dropped to 41.3% for the MSBooster-removed peptides. This further supports the idea that peptides gained with DL-based rescoring in MSBooster are more reliable than those that are removed.

An important feature of MSBooster is its ability to handle peptides with post-translational modifications (PTMs) that are not predicted by the DL spectral prediction model. In DIA-NN v1.8, cysteine carbamidomethylation, methionine oxidation, N-terminal acetylation, phosphorylation, and ubiquitination are supported. A multitude of other PTMs exist; in the FragPipe HLA workflow, for example, cysteinylation is an important PTM to consider, as it plays a role in T cell recognition [61]. Rather than precluding the inclusion of other PTMs in the search or rescoring steps, MSBooster obtains the predicted spectrum for the unmodified peptide and shifts the m/z values of the PTM-containing fragments while retaining their predicted intensities. RT values are the same as those of their peptide counterparts, excluding the new PTMs (e.g., a jointly biotinylated and phosphorylated peptide will use the RT of the phosphorylated peptide). To explore how fragment peak shifting affects the results, we examined the distributions of spectral and RT feature scores for accepted PSMs after Philosopher filtering (**Supplementary Fig 4**). Each group of PSMs contained only the PTM listed (i.e., the PSMs in the carbamidomethylated C group were matched to peptides that only contained that PTM, and no oxidized M, etc.). As expected, unmodified, carbamidomethylated C, and oxidized M PSMs had high spectral similarities and low RT differences, as DIA-NN included them in the training set. Interestingly, although acetylated N-terminal peptides were in the training set, their spectral similarity score distributions were lower than those of the other peptides. Cysteinylation had a similar distribution of PTMs on which DIA-NN was trained. This could mean that cysteinylation does not have a major impact on fragment intensity or that cysteinylated PSMs with lower scores were excluded after FDR filtering. PSMs with pyro-glutamation events from Q had the worst distribution of the PTMs considered. Although this PTM noticeably changes the fragment ion intensities and RTs (the RT shift in pyro-Glu peptides has been previously recapitulated [62]), they may still pass the FDR threshold because of the high discrimination power of other non-DL features. Overall, our analysis shows that MSBooster allows re-scoring of any PTM-containing peptide, even if a small penalty is applied in the case of certain PTMs (not yet supported by DIA-NN prediction module).

### Direct identification form DIA data

The DIA strategy offers the benefit of monitoring all precursors (within the specified mass range, e.g. 400-1200 Da) and their fragments across the retention time, thereby avoiding the stochasticity of DDA which can only produce MS/MS scans for a limited number of precursors. We extended MSBooster to rescoring of peptide identifications from DIA data. In FragPipe, peptide identification from DIA data can be performed in two ways: (1) with MSFragger-DIA, which identifies peptides from DIA MS/MS scans by direct database searching; and (2) by first processing the DIA MS files using DIA-Umpire [7] to extract pseudo MS/MS spectra, followed by searching with MSFragger as regular DDA data. We tested both approaches on a dataset of six melanoma cell lines (analyzed in duplicate; 12 files in total) taken from [63]. Using MSFragger-DIA, DL-based rescoring with MSBooster using both spectral and RT features increased peptide and protein identifications by 16.6% and 8.9%, respectively (**Fig 3a, c**). Using the DIA-Umpire based workflow, the number of peptide and protein identifications increased by 16.6% and 9.0%, respectively (**Fig 3b, d**).

**Figure 3.**
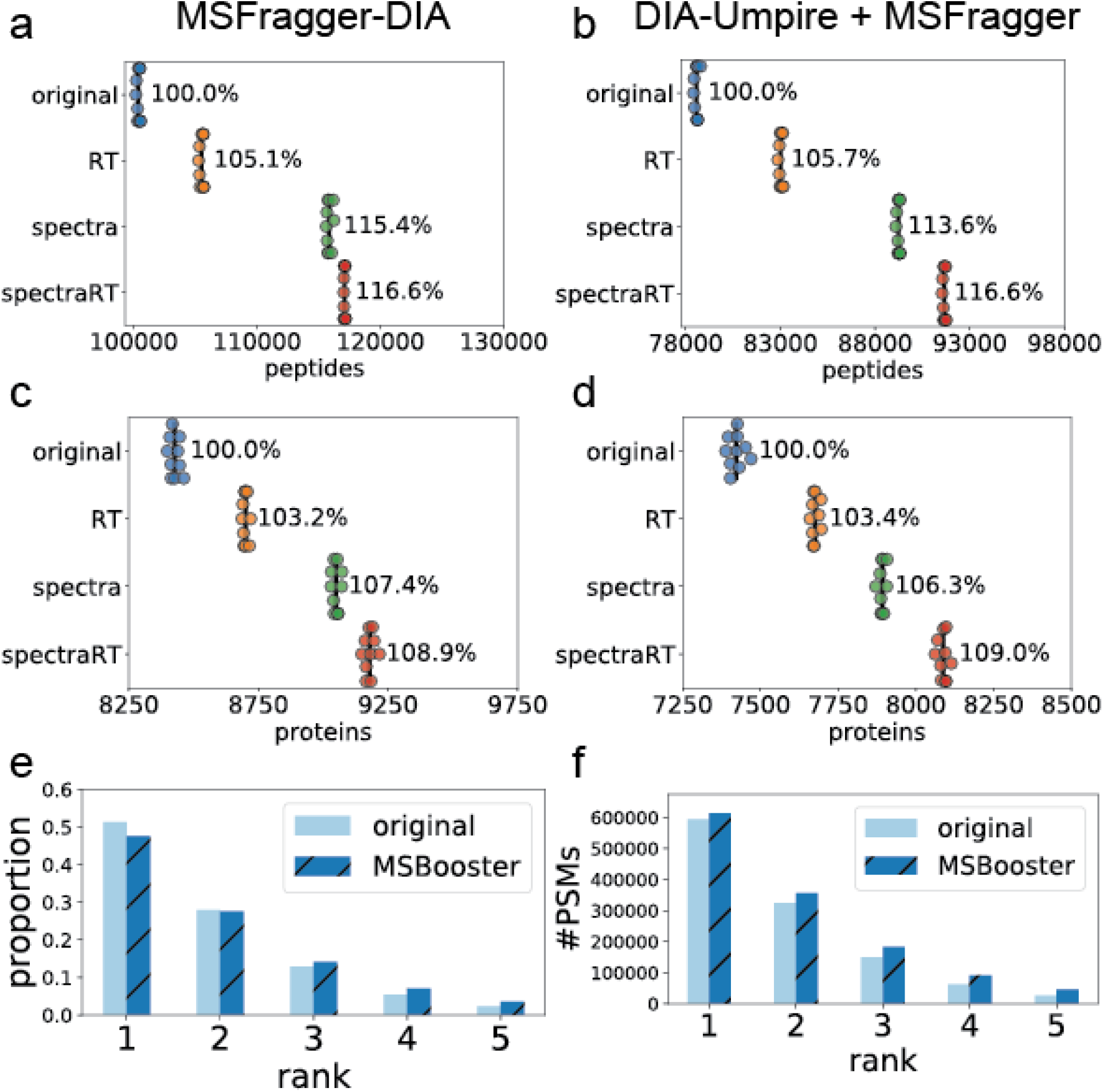
DIA dataset. (a-d) Swarmplots of the number of peptides (a-b) or proteins (c-d) reported at 1% FDR when using MSFragger-DIA (a, c) and DIA-Umpire with MSFragger (b, d). The proportion of PSMs (e) or total PSMs (f) from each of the five ranks reported by MSFragger-DIA. The darker, diagonally dashed bars represent results after spectral and RT rescoring, while the lighter, solid bars represent the results without using deep learning features.

The benefit of rescoring MSFragger-DIA results with MSBooster applies not only to the top scoring, but also to lower-ranking PSMs. By default, MSFragger-DIA reports up to 5 PSMs per MS/MS scan. While the initial MSFragger rankings were based on hyperscore, other features provided orthogonal information (**Supplementary Fig 2b**) and helped rescue true PSMs with lower hyperscores. With MSBooster, while the total number of PSMs passing the 1% FDR increases across all ranks, a higher proportion of accepted PSMs are from ranks 3 and below, while the relative proportion from ranks 1 and 2 decreases (**Fig 3e-f**). Note that MSFragger provides a rank feature for Percolator to use, but this feature rarely had a Percolator weight above 0.1; for comparison, the hyperscore often had a weight above 2. This indicates that Percolator gives little preference to higher-ranking PSMs in DIA data, allowing multiple PSMs with high spectral and RT similarities to be identified for a single DIA MS/MS spectrum, reflecting the multiplex nature of DIA MS/MS scans.

### Single cell proteomics

Single-cell proteomics provides a view of the proteomes of individual cells. The lower level of maturity of technological platforms, along with the increased stochasticity of peptide identification due to cell-to-cell variability, make single cell proteomics another promising area for DL-based PSM rescoring. We tested MSBooster on single cell data from the nanoPOTS platform [64] generated using an Orbitrap Fusion Lumos Tribrid instrument (Thermo Fisher scientific). Briefly, we analyzed the data obtained from 1, 3, 10, or 50 cells. Single-cell MS/MS spectra differ from bulk-cell spectra in terms of the number of fragments matched and the degree of fragment ion intensity suppression [65]. When looking at the scores of top target PSMs with an increasing number of cells in the sample (from 1 to 50), we found a trend (**Fig 4a**) that with more cells there was a gradual increase in the median spectral similarity among confidently identified target PSMs (i.e., PSMs with expectation values lower than the lowest decoy PSM expectation value, see **Methods**). As a reference, bulk secretome data obtained from [66], also generated on an Orbitrap Fusion Lumos instrument, demonstrated a significantly higher median spectral similarity score. With respect to RT values, there was a decrease in the median RT difference between 1 and 3 cells, due to an insufficient number of PSMs for optimal RT calibration in MSBooster with one cell only (**Fig 4b**). However, the median RT difference did not decrease past 3 cells, because the RT difference should not change once there is a sufficient number of PSMs for RT calibration. The bulk cell RT score distribution was excluded from the comparison, because the RT score depends on the individual LC set up. Surprisingly, when looking at the different cell numbers separately, we did not notice a clear relationship between the cell number and MSBooster performance (**Fig 4c-d**). The single-cell (1 cell) experiment gained 4.7% and 2.8% peptide and proteins, respectively, with spectral and RT rescoring. The few-cell data (3, 10, and 50 cells) gained up to 10.6% peptides (in the 50-cell dataset) and 10.6% proteins (in the 3-cell dataset). The RT feature outperformed the spectral feature in many instances.

**Figure 4.**
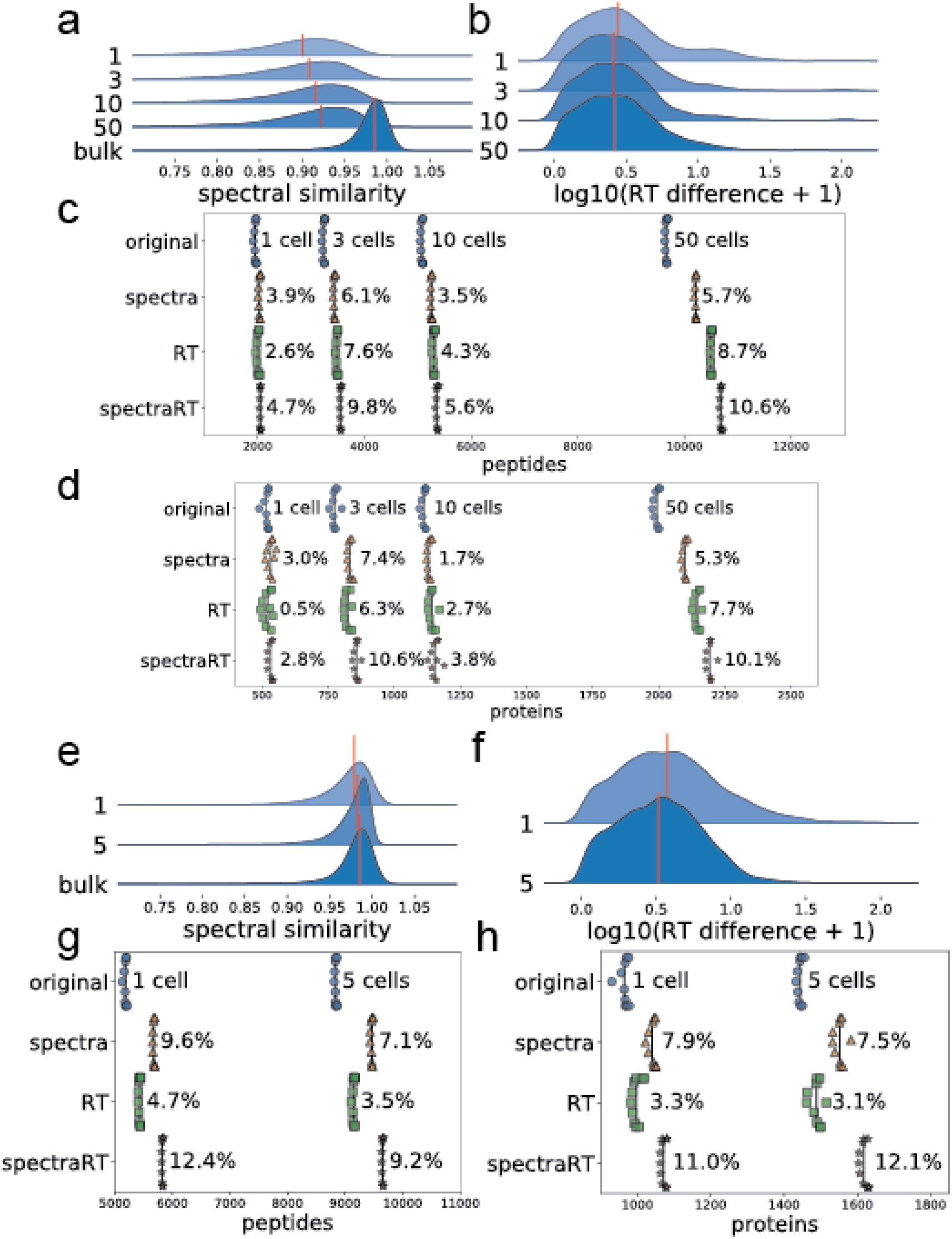
Single cell proteomics data. (a-d) Results for nanoPOTS data. (a-b) Ridge plots showing the distribution of the spectral (a) and RT (b) feature scores of confident target PSMs for different numbers of cells. The red line indicates the median value. The bulk cell sample is from PXD026436. The RT feature was log normalized for better visualization. (c-d) Swarmplots of the number of reported peptides (c) and proteins (d) when using different features for Percolator rescoring. (e-h) are the same as (a-d), but for the DISCO single cell dataset.

Because single-cell proteomics methods are being rapidly developed and modified, we tested another dataset produced on a Q-Exactive MS with a different sample processing protocol (DISCO) [46]. Data from 1 and 5 cells were available. While we see similar trends of increasing median spectral similarity and decreasing median RT difference with an increasing number of cells, median spectral similarity is already above 0.95 for single cells in these data (**Fig. 4e-f**). In comparison, even 50 cells in the nanoPOTS data had a median spectral similarity below 0.95 (**Fig 4a**). We also noted significant differences in the decoy PSMs’ spectral similarity distributions (median of 0.49 vs 0.31, nanoPOTS vs DISCO) and differing numbers of PSMs reported per replicate (mean of 2924 vs 19878, nanoPOTS vs DISCO). We can see the effect of the higher similarity reflected in the number of peptide and protein identifications achieved with rescoring (**Fig 4 g-h**). In this dataset, the spectral similarity feature always outperformed the RT feature. Using both features, MSBooster provides a 10-15% boost.

### timsTOF PASEF data with ion mobility separation

Next, we evaluated the performance of MSBooster on data from a standard proteomics sample (HeLa tryptic digest standard) analyzed using Parallel Accumulation–Serial Fragmentation (PASEF) on a timsTOF Pro mass spectrometer [2]. Using both spectral and RT features, we achieved a 3.9% and 2.7% increases in peptide and protein identification, respectively (**Fig 5a-b**). While this seems to be a minor increase, it highlights that FragPipe’s default workflow for conventional tryptic searches performs well even without DL-based rescoring. In this case, MSFragger with just a hypersore can find most peptides and proteins without the help of the other features reported by MSFragger and without DL-based features (**Fig 5a-b**).

**Figure 5.**
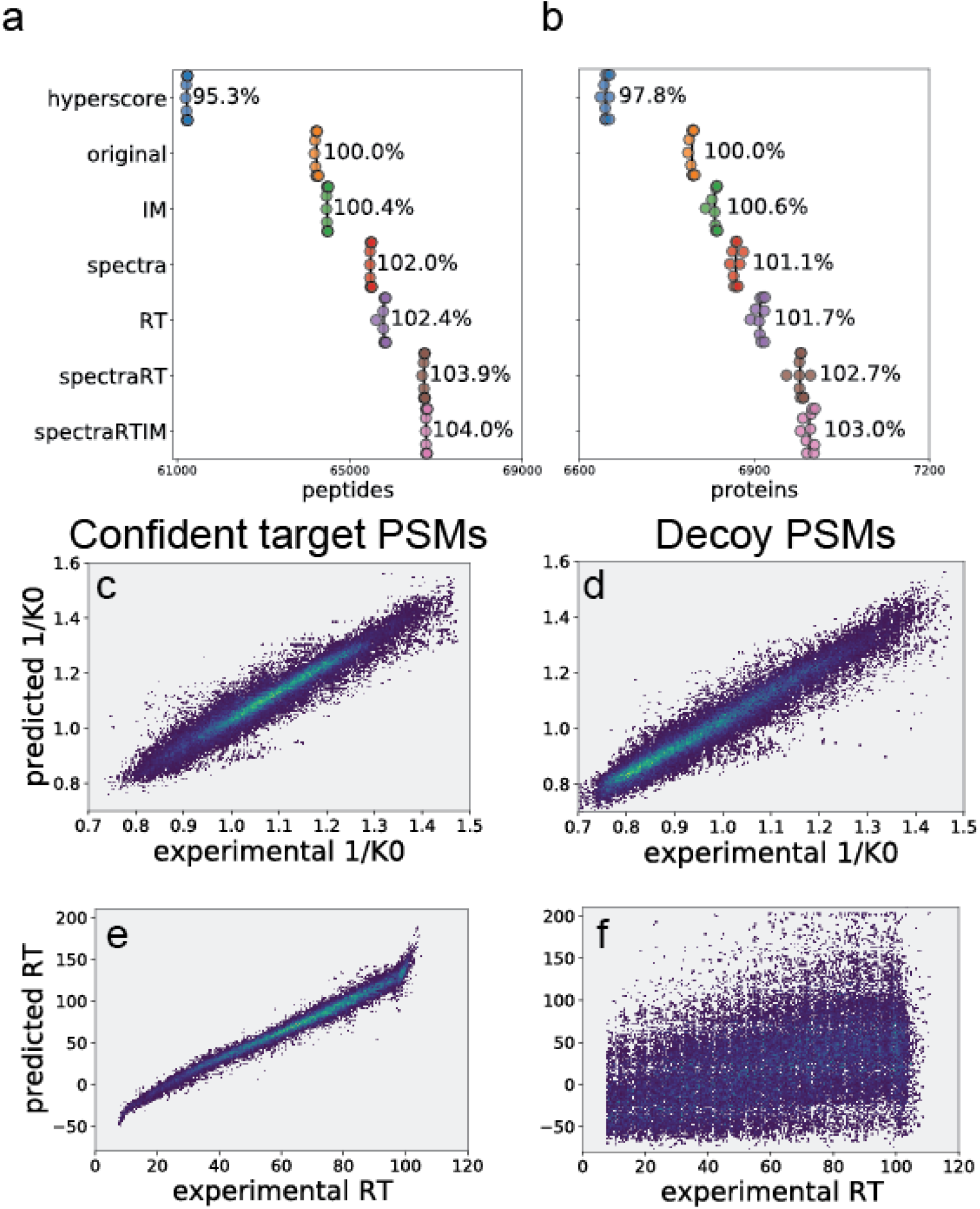
timsTOF PASEF HeLa dataset. (a-b) Swarmplot of peptides (a) and proteins (b) reported at 1% FDR. (c-f) Scatter density plots showing the relationships between DIA-NN predicted and experimental IM (c-d) and RT (e-f) values in seconds for peptides with charges 2 and above. Confident target PSMs are shown in (c) and (e), decoy PSMs in (d) and (f). The brighter colors correspond to higher densities of PSMs.

Ion mobility is an additional method for separating precursors prior to MS/MS sequencing. As such, DL models have been extended to predict ion mobility or related collisional cross section values [39, 44]. To assess the utility of predicted IM for PSM rescoring, we ran MSBooster with IM features analogous to MSBooster’s RT features (see **Methods**). However, we observed a negligible increase in the number of identified peptides and proteins, below 0.5% (**Fig 5a-b**), with the addition of the IM score. Furthermore, IM features barely had sufficient discriminative power to separate targets from decoy PSMs at 5% FDR. The weakness of the IM features may be explained by the high dependence of the IM on the precursor mass and charge. Because decoy PSMs still have the same charge and similar mass as the unknown true target precursor, predictions of their 1/K0 values are still highly correlated with the experimental value (**Fig. 5c-d; Supplemental Fig 5**). There is not as much of a spread for IM as for RT prediction (**Fig. 5e-f**). Overall, while the target PSMs showed a slightly different distribution of IM feature scores than the decoy PSMs (**Supplementary Fig 6e**), it was not sufficient to warrant their use in MSBooster.

We also noted a higher density of decoy PSMs at lower 1/K0 values (around 0.7-0.8) compared to confident target PSMs (**Fig 5c-d**). This preference for lower 1/K0 values among decoy PSMs persists for each individual charge state between 1 and 3 but starts to disappear for charge 4 and above. Similarly, decoy PSMs tend to have lower masses and shorter lengths than target PSMs for charges 1-3 (**Supplementary Fig 7**). The higher rate of decoy PSMs at lower inverse IMs/masses/lengths may be attributed to fewer fragments being produced by shorter peptides. The peptide mass and length have already been incorporated as features in the MSFragger-generated pin file. We tested adding 1/K0 values (not to be confused with the DL-based IM features in MSBooster) as a feature for Percolator as well, but again observed a negligible impact (**Supplementary Fig. 5**).

### Multiple correlated features

Because MSBooster can calculate several variants of spectral, RT, and IM features, we evaluated whether there was value in using multiple correlated features for PSM rescoring (**Supplementary Fig 8**; all tested features are described in the **Supplemental Note**). This idea was spurred by the finding that even correlated features may not be truly redundant and may work well to provide better separation between classes [67]. For all datasets, we annotated Percolator input files with either single features (“spectraRT” and “spectraRTIM”) or all available features listed in the **Supplemental Note** (“multipleSpectraRT” and “multipleSpectraRTIM”). In most analyses, the use of multiple correlated features resulted in a minor (<1%) increase in identification numbers. Occasionally, the numbers decreased by an equally small amount (**Supplementary Fig 8**). To investigate this further, we returned to the HLA immunopeptidome dataset. With the use of multiple correlated features, compared to using only a single feature for each feature type (as in **Fig. 2**), 165 peptides were lost, and 184 were gained. Both groups of lost and gained HLA peptides had comparable binding rates (123/165 vs 134/184, p > 0.05, chi-square test) and similar sequence motifs (**Supplementary Fig 8**). However, because it is difficult to rule out all scenarios where using correlated features may be beneficial, we provide an option for FragPipe users to enable their use.

## Discussion

MSBooster is a new addition to the FragPipe computational platform that provides a boost in the number of identified PSM, peptides, and proteins by generating deep learning-based features for PSM rescoring with Percolator. It automatically runs a model (DIA-NN) to acquire predictions for different physicochemical properties (RT, IM), and MS/MS spectra, and generates features based on these predictions to expand the list of scores for each PSM that are useful for discriminating between true and false matches. We evaluated the improvements provided by MSBooster in various experimental workflows available in FragPipe (HLA immunopeptidomics, timsTOF PASEF data, single cell proteomics, and DIA data). We observed robust gains across all applications, especially in analyses exhibiting a large “grey zone” in the true vs. false PSM separation space, such as HLA immunopeptidomics and direct identification from DIA data searches. A notable benefit of MSBooster is that the whole peptide library does not need to be predicted; only high-scoring candidates identified by MSFragger are evaluated using MSBooster, saving a large amount of time, especially for nonspecific searches. Furthermore, although not fully explored in this work, MSBooster can be used to process the MSFragger output for DDA data with multiple PSMs reported per spectrum, potentially assisting with the identification of co-fragmenting precursors present in DDA data [68].

Using the HLA immunopeptidome dataset, we also examined MSBooster’s feature score distributions for peptides with different PTMs that were allowed in the search and provided an experiment-level view of how MS/MS spectra/RT for these modified peptides deviated from their unmodified counterparts. In single-cell proteomics data, we observed that although it is possible to achieve improvements of nearly 15%, successful PSM rescoring is contingent upon our ability to accurately predict MS/MS spectra in such studies. A recent study revealed that single-cell and bulk cell proteomics MS/MS spectra differ [65]. Models trained specifically on single-cell spectra or bulk cell models that have been fine-tuned [40] on single-cell spectra may be better suited to this rescoring task.

We observed a marginal impact of adding IM features when analyzing standard HeLa tryptic digest timsTOF PASEF data. However, this does not mean that ion mobility added information will not be useful in other scenarios. IM may improve the resolution of peptidoforms with isobaric modifications (e.g., in glycopeptide identification workflows [69]) or assist with PTM site localization [70, 71]. The minimal strength of IM features observed in this work may also be due to the limitations of a linear SVM model in Percolator. Thus, more flexible models [72] for PSM rescoring could be investigated.

Future work will focus on adding more flexibility to MSBooster with respect to which prediction model is used to generate the predicted spectra. For example, PredFull [73] is a full spectrum prediction model that predicts intensities for every m/z bin, rather than predicting specific ion types such as y- and b-ions. Therefore, it may be able to report internal fragment ion intensities, which could provide more rescoring information for HLA peptides lacking the basic C-terminal residues - common to tryptic peptides – that help to create a strong y-ion series. Other scenarios where non-y/b ions are abundant and may also benefit from using PredFull is rescoring ETD MS/MS spectra, or spectra produced by peptides with PTMs that incur neutral losses. Support for diverse prediction models within MSBooster will be particularly useful for studying PTMs, where models such as pDeep2 [42], MS2PIP [37], and DeepLC [74] are expected to perform better than simply shifting fragment ions or using the same RT for both modified and unmodified peptides. The increased use of labeled quantification with tandem mass tags (TMT) has also allowed DL models to be trained for this purpose [75]. Support for more DL models may also benefit HLA rescoring, where models such as Prosit have been trained on nonspecific peptides [47]. Finally, transfer learning and fine-tuning implemented in pDeep3[40] and AlphaPeptDeep [41] may help to analyze MS/MS spectra acquired using different fragmentation mechanisms, or when identifying peptides containing rare PTMs.

## Methods

### MSBooster workflow

Workflows with and without MSBooster are depicted in **Fig 1**. In DDA experiments, MS/MS spectra are searched using MSFragger. For peptide identification from DIA data, either full MS/MS spectra are searched using MSFragger-DIA, or DIA-Umpire extracted pseudo-MS/MS spectra are searches using MSFragger as conventional DDA files. MSFragger produces pepXML and ‘pin’ files, and the latter are used as input into MSBooster. The pepXML files are not used by MSBooster but are necessary for converting the Percolator output files into pepXML files for subsequent protein inference analysis using ProteinProphet. To obtain a list of peptides for DL model prediction, MSBooster iterates through all pin files to obtain all target and decoy peptides matched to at least one PSM. Peptides with the same sequence but different PTMs and/or charges are treated as different peptides. This list is then passed to DIA-NN, which is available as part of FragPipe, to obtain a prediction file. This strategy is significantly faster than predicting the entire in-silico digested proteome, as only peptides reported as potential hits by MSFragger (top ranking peptides per spectrum) are submitted for prediction. Peptides with PTM(s) not supported by the DIA-NN model rely on predictions for the peptide without the unsupported PTM(s). MSBooster adds a shift in fragment m/z to accommodate the new PTMs, but the fragment intensities remain the same.

The core of MSBooster is the feature calculation step. First, the predictions from DIA-NN are loaded. Then pairs of mzML (or MGF) and pin files are sequentially loaded for processing and DL-extended PSM feature table generation. For DIA data, because multiple peptides may contribute to a single MS/MS scan, the experimental spectra are revised after each PSM has its features calculated. Borrowing from MSFragger-DIA, the highest intensity experimental MS/MS peak within the fragment error tolerance of the reported predicted fragment is removed from the experimental spectrum. The predicted spectra of lower-ranking PSM peptides can no longer have their fragments matched to these removed peaks, because using the same MS/MS peaks for multiple PSMs from the same scan can lead to spurious hits. Once all PSMs from a single pin file are loaded, RT and IM calibrations are performed. In the final step, MSBooster iterates through the pin file row by row, and calculates and adds the desired features. This process is repeated until all pin files have DL features calculated and added. The list of all available features is described in the **Supplemental Note**.

### Determination of the best features

Several metrics exist for calculating the similarities between experimental and predicted spectra. Although cosine similarity is commonly used, several features were tested to determine which metrics could provide the greatest gains in the identification numbers (**Supplementary Fig 10**). Percolator is non-deterministic because of the random splitting of PSMs for training and testing, which can be controlled with a random seed. Thus, Percolator was run ten times for each feature, and the number of peptides reported after Philosopher filtering was counted. For spectral similarity features, the greatest boosts were consistently obtained with unweighted spectral entropy [76]. For the RT features, “delta RT loess” tended to do the best. Interestingly, “delta RT loess normalized” performed better when there were a small number of cells in the nanoPOTS data [64] (**Supplementary Fig 10e-h**). For the IM features, the IM probability feature performed the best. These single features are used for rescoring unless otherwise noted. The distributions of each score for all PSMs, targets and decoys, in the different datasets are shown in **Supplementary Fig 6**.

### MSFragger search and FDR control

Database searches were performed using MSFragger v3.4 in FragPipe v17.2 with Philosopher v4.1.1. The workflows used for each dataset are as follows: HLA immunopeptidome [56] (*nonspecific-HLA-C57* workflow), melanoma DIA data [63] with MSFragger-DIA (*DIA_SpecLib_Quant)* and with DIA-Umpire (*DIA_DIA-Umpire_SpecLib_Quant*), HeLa timsTOF [2] (*Default*), single cell proteomics with nanoPOTS [64] (*Default*), single cell proteomics with DISCO [46] (*Default*), and secretome [66] (*Default*). The HLA workflow was revised to add “--mods M:15.9949” to the Philosopher filter to perform group-specific FDR estimation [77] using the following three categories: unmodified peptides, peptides with oxidized M only, and peptides with any other modification. The nanoPOTS data were analyzed in separate experiments based on the number of cells (1, 3, 10, or 50).

### Deep learning predictions

MSBooster can be used with any RT/IM/spectral prediction tool. In this work, we chose DIA-NN [43, 44] because of its speed and ease of execution within FragPipe. DIA-NN v1.8 was used to predict RT, IM, and MS/MS spectra for the top 12 most intense singly and doubly charged b- and y-ions. Predictions were made for each unique combination of peptide sequence, modifications, and charge. DIA-NN v1.8 supports the predictions for peptides with carbamidomethylated cysteine, oxidized methionine, N-terminal acetylation, phosphorylation, and ubiquitination. For other PTMs such as pyro-glutamation, DIA-NN did not adjust MS/MS fragment peak intensities, but MSBooster shifted the peaks to the appropriate m/z values. The RTs and IM values for peptides with unsupported PTMs remained the same as for counterparts without the PTM.

### Spectral similarity calculation

To calculate the spectral similarity, the highest intensity fragment ions within the m/z error tolerance of the predicted fragment ions are obtained. Therefore, similarity calculations are performed using vectors of the same length. Predicted and matched fragment ions from the experimental MS/MS spectra were normalized before similarity calculation (see **Supplemental Note**).

### Retention time and ion mobility calibration

Local regression (LOESS) is used to calibrate the experimental to the predicted RT and IM values, followed by monotonic regression. A different ion mobility model is trained for each charge. To train the regression models, a subset of PSMs with expectation values below a preset threshold is used. If fewer than 50 target PSMs with sufficiently low expectation values are available, linear regression is performed. For both DDA and DIA, only rank 1 PSMs are considered for the regression. To calculate the difference between the predicted and experimental RT, the experimental RT is first calibrated to the predicted scale using the regression model, followed by calculating the difference between the calibrated RT for that MS/MS scan and the predicted RT for that peptide. The same is performed for the IM.

### Kernel density estimation of predicted retention time and ion mobility distributions

Empirical distributions of predicted RT/IM values were generated using Statistical Machine Intelligence and Learning Engine (Smile) implementation of kernel density estimation (KDE) with a Gaussian kernel (https://haifengl.github.io/api/java/smile/stat/distribution/KernelDensity.html). The following discussion uses RT, but the same applies to IM. The empirical range is divided into bins. For each PSM, the predicted RT is placed into a bin determined by the experimentally observed RT. The number of times its RT value is added to the bin is weighted by its expectation value; that is, a higher-confidence PSM with a low expectation value will have its predicted RT added to the bin more times than a lower-confidence PSM with a high expectation value. After all predicted RTs are placed in their respective bins, KDE is used to generate empirical distributions. These distributions can be used to estimate the probability of having a PSM with a predicted RT value given its experimental RT and are not subject to the monotonic constraint of the LOESS model. The same procedure is applied to the IM to generate distributions of the predicted IM, separating the PSMs by charge state. The features from MSBooster that use these probabilities also add a uniform prior distribution to the KDE-generated distribution. This uniform prior helps to dampen the effects of bins with fewer entries. For example, if an experimental RT bin contains a single PSM, not using a uniform prior would result in an artificially high probability for that PSM.

### HLA motif analysis

Before using HLA peptides for analysis, they were filtered to be between lengths 7 and 12. Position weight matrices were generated using GibbsCluster 2.0 [58] (https://services.healthtech.dtu.dk/service.php?GibbsCluster-2.0). The binding affinity of the peptides to the A*02:01 MHC was determined using NetMHC 4.0 [60] (https://services.healthtech.dtu.dk/service.php?NetMHC-4.0) using the default settings.

### Statistical analysis and figure generation

Statistics and figures were generated in Jupiter Notebooks using Python 3.7.6. The scatter density plots for Figure 5 and Supplemental Figure 5 require a separate Anaconda environment with Python 3.8.3.

### Hardware

FragPipe was run and timed using Java 16.0.1. A command-line version of MSBooster was run on a Windows desktop with 12 logical CPU cores (Intel(R) Core™ i7-8700 CPU @ 3.20 GHz) and 32 GB of memory. This was essential for automating the testing with different MSBooster features.

## Data availability

MS/MS datasets used in this study can be found at the ProteomeXchange Consortium and the PRIDE partner repository (PXD)[78] or at the MassIVE repository (MSV).

- HeLa timsTOF DDA: PXD010012 [2].
- HLA peptidome: MSV000087743 [56].
- Melanoma DIA: PXD022992 [63].
- Single cell nanoPOTS: MSV000085230 [64].
- Single cell DISCO: PXD019958 [46].
- Secretome: PXD026436 [66].

All MSFragger produced pepXML files and MSBooster-annotated pin files are available at https://doi.org/10.5281/zenodo.7072124

## Code availability

MSBooster code is available freely and as open source at https://github.com/Nesvilab/MSBooster.

## Author Contributions

K.L.Y. and A.I.N. conceived the study. K.L.Y. developed the algorithm, wrote the software, and analyzed the results. F.Y. assisted with the algorithm and software development. F.Y. and G.C.T. provided support for integration of MSBooster in FragPipe. V.D. and M.R. provided resources pertaining to DIA-NN. K.L.Y. and A.I.N. wrote the manuscript. A.I.N. supervised the study.

