## Supplementary Note for "MSBooster: Improving Peptide Identification Rates using Deep Learning-Based Features"

**Supplemental Note**

*Supplemental Note 1. Features*

Available features in MSBooster are listed below, along with an equation and source for less commonly used metrics. By default in FragPipe, the unweighted spectral entropy and delta RT LOESS only are used.

- Spectral similarity: Unweighted spectral entropy requires fragment ion intensity vectors to sum to 1. Other metrics require unit normalization (i.e. the sum of squared intensities equals 1). p stands for predicted fragment, P for predicted intensity vector, m for matched experimental fragment, and M for matched experimental intensity vector. By default, the top 12 highest predicted intensity fragments are used.
    - Bray Curtis[1]: $1 - \frac{\Sigma_{i=1}^{n} |p_i - m_i|}{\Sigma_{i=1}^{n} p_i + m_i}$
    - Pearson's correlation: If no matched experimental fragment ions, value is -1
    - Dot Product
    - Unweighted spectral entropy[2]: $1 - \frac{2S_{PM} - S_P - S_M}{ln\,4}$, where entropy S = $-\sum_{i=1}^{n} f_i \, ln \, f_i$, and $S_{PM}$ is the sum of predicted and matched vectors $S_P$ and $S_M$ divided by 2
- Retention time similarity
    - delta RT loess: A monotonic local regression curve is fitted to each run using some subset of the PSMs with lowest expectation values. After calibrating experimental RTs to the predicted RT scale, the difference between calibrated RT and predicted RT for a peptide is reported.
    - delta RT loess normalized: Similar to delta RT loess but divided by the interquartile range of predicted RTs for high confidence PSMs in the experimental RT vicinity.

    - RT probability Unif Prior: The experimental RT range is binned. For each bin, kernel density estimation with a Gaussian kernel is performed on the high confidence PSMs' predicted RTs. A PSM can be rescored by matching its experimental RT to the appropriate bin and calculating the probability from the empirical distribution.
- Ion mobility similarity
    - Features are like those for RT but done separately for each charge. Only charges +1-7 are supported.


1 Toprak, U. H. *et al.* Conserved Peptide Fragmentation as a Benchmarking Tool for Mass Spectrometers and a Discriminating Feature for Targeted Proteomics. *Molecular & Cellular Proteomics : MCP* **13**, 2056-2056 (2014). https://doi.org:10.1074/MCP.O113.036475

2 Li, Y. *et al.* Spectral entropy outperforms MS/MS dot product similarity for small-molecule compound identification. *Nature Methods* **18**, 1524-1531 (2021). https://doi.org:10.1038/s41592-021-01331-z
