## Supplementary Figures for "MSBooster: Improving Peptide Identification Rates using Deep Learning-Based Features"

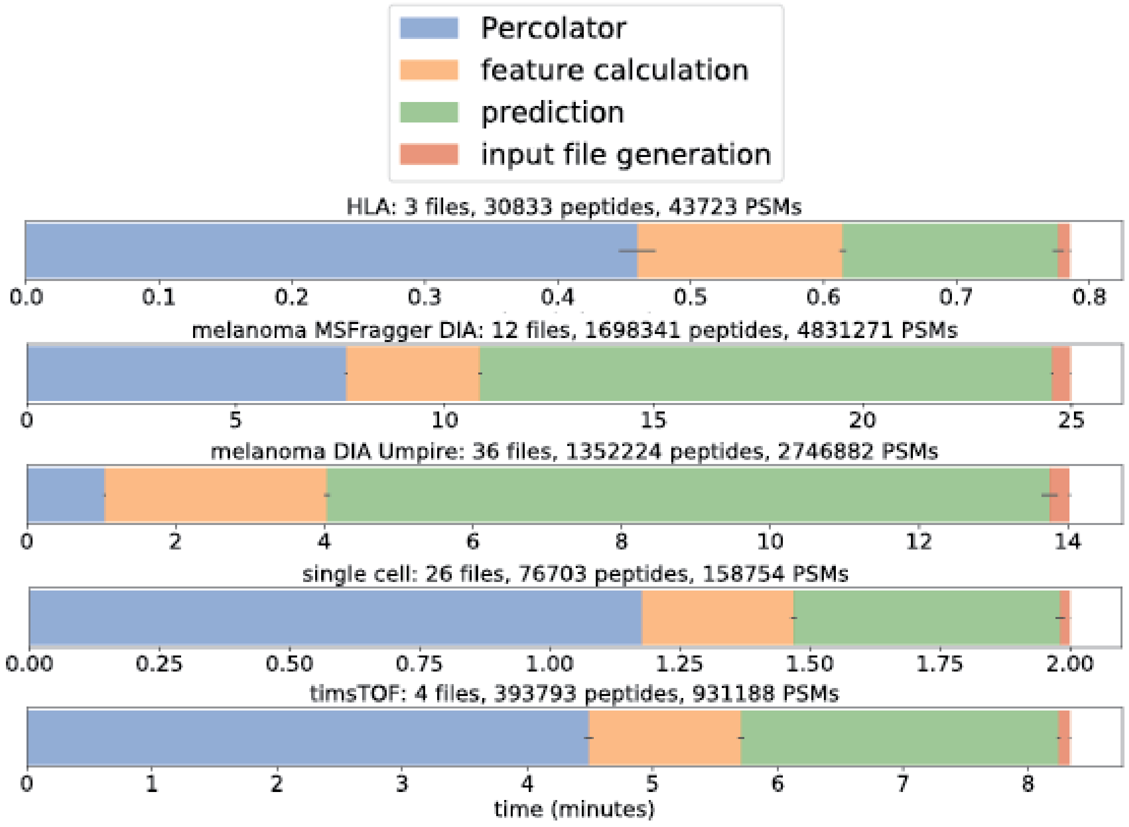


**Supplemental Figure 1.** Timing of MSBooster steps and Percolator rescoring. Error bars show the standard deviation from running the software ten times. For each subfigure, the dataset, the number of mzML/pin file pairs, the number of unique peptides in the pin files, and the total number of PSMs are listed.

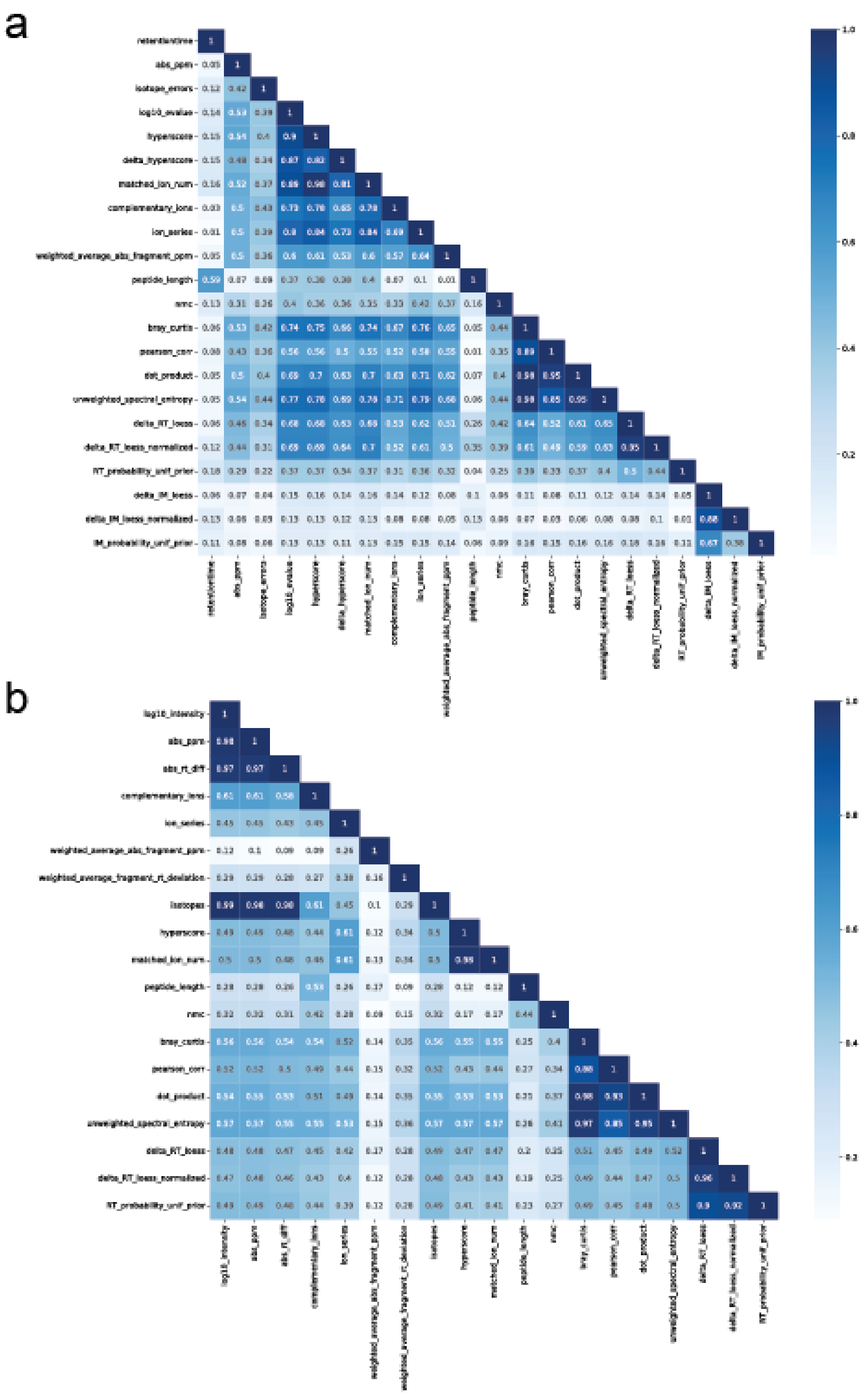


**Supplemental Figure 2.** Feature correlations. Spearman's correlation was calculated between MSBooster features and all features reported by MSFragger (a) and MSFragger-DIA (b). (a) was produced from one timsTOF file, (b) from one melanoma file.

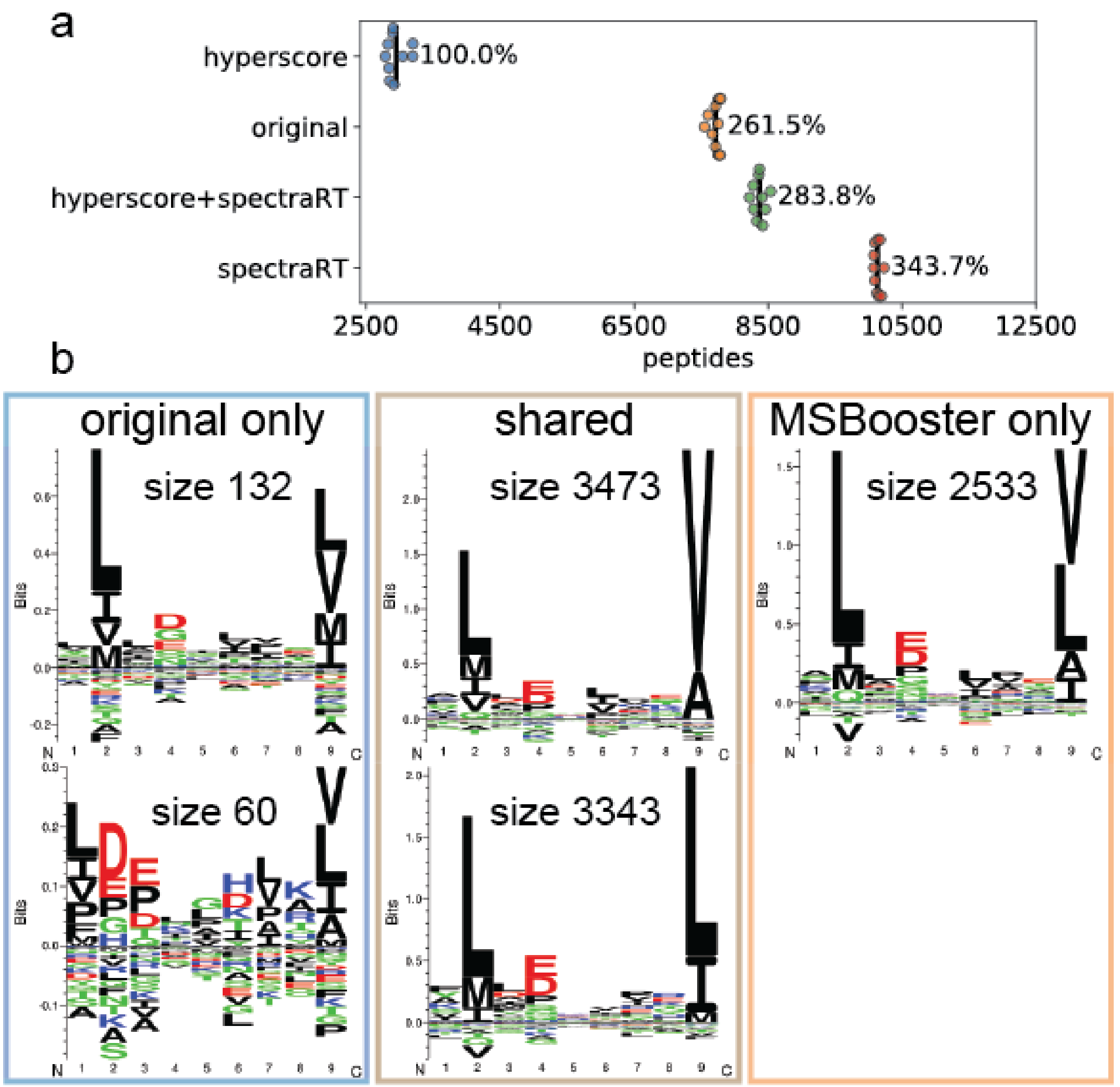


**Supplemental Figure 3.** Utility of non-deep learning features from MSFragger. (a) Swarmplot of the number of HLA peptides reported when only using hyperscore or hyperscore and the deep learning features without other MSFragger features (hyperscore+spectraRT). (b) GibbsCluster-generated motifs assigned to each peptide subset from the Venn diagram in Fig 2b.

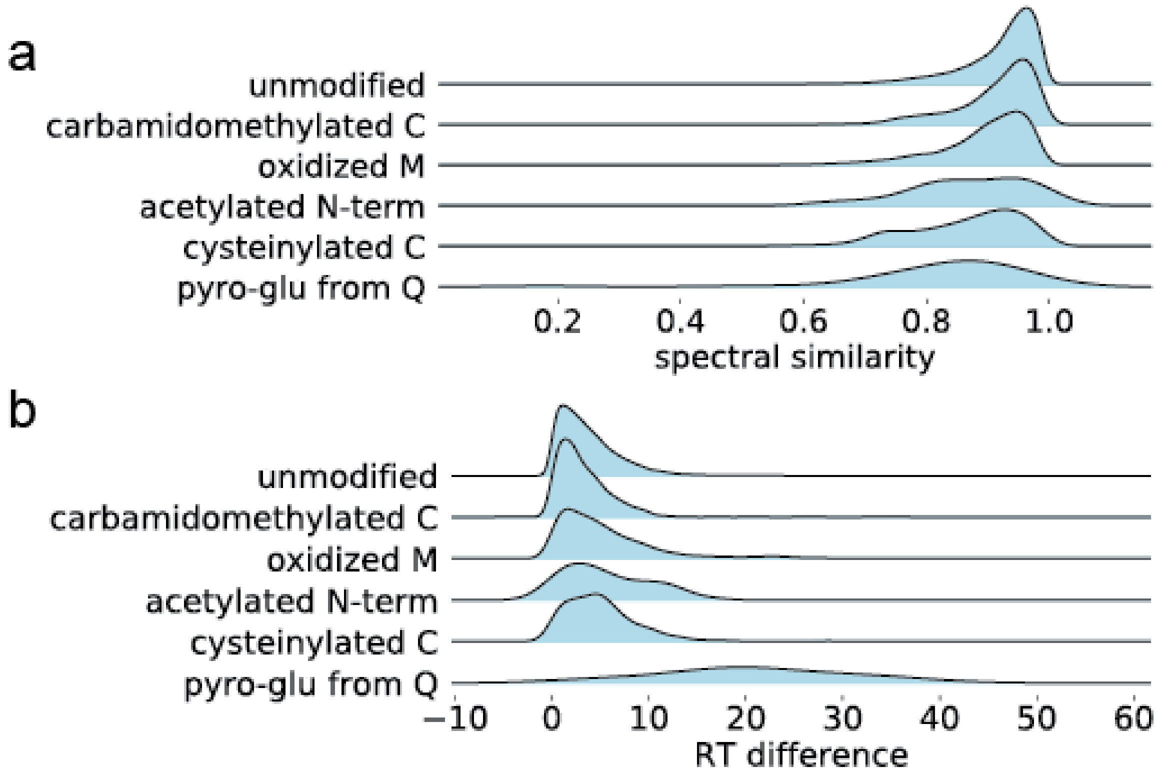


**Supplemental Figure 4.** Score distributions for unmodified and modified HLA peptides. (a) Spectral similarity values for accepted PSMs at PSM FDR < 1% and peptide FDR < 1%. (b) The same as (a) but for RT difference values. Note that these values are not log-normalized as in other figures. Pyro-glutamation on E was included in the search, but only one was detected and therefore excluded from visualization.

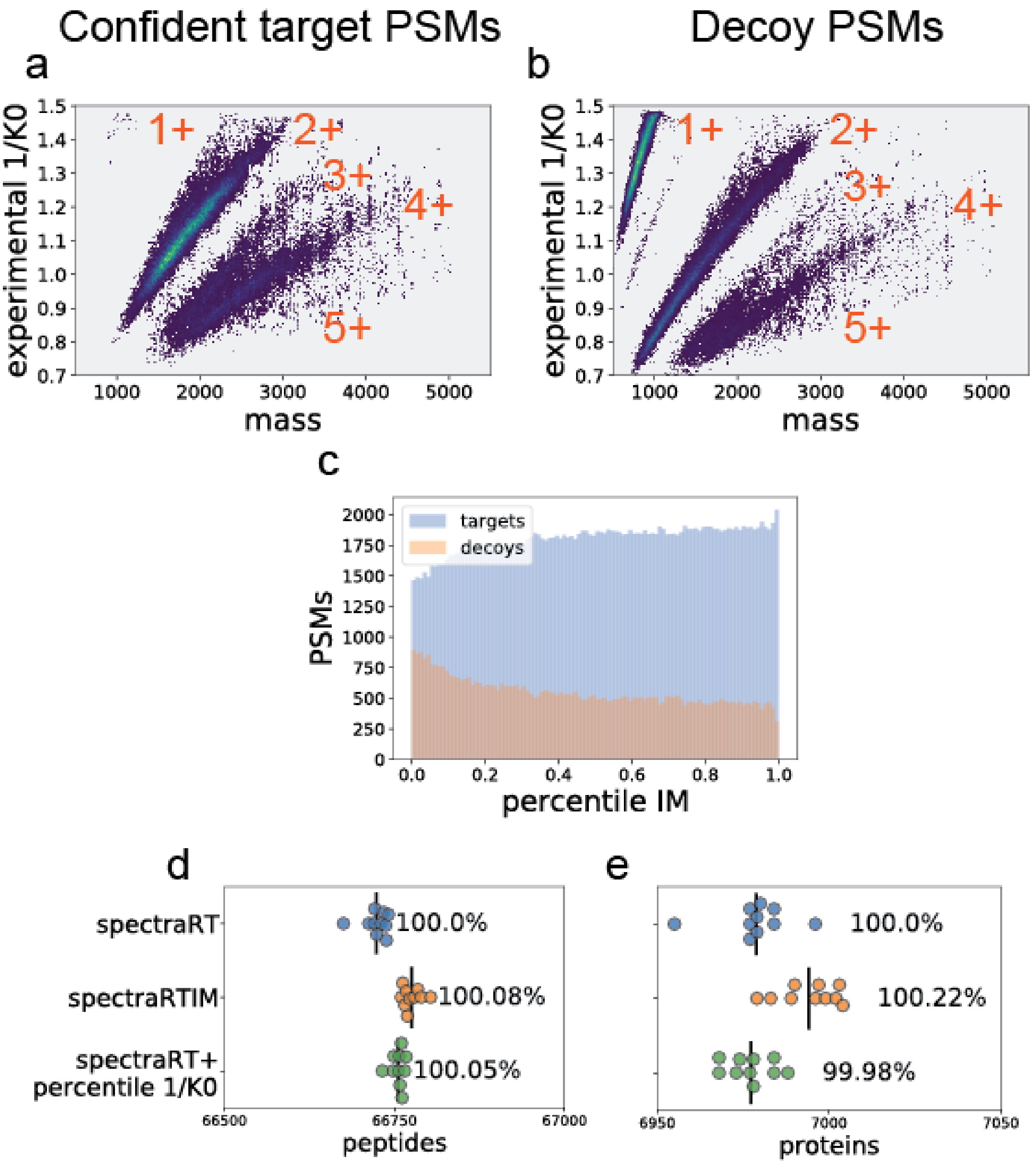


**Supplemental Figure 5.** Inverse ion mobility dependence 1/K0 on mass and charge. (a-b) Scatter density plots are shown for (a) confident target PSMs and (b) decoy PSMs. (c) Histogram distributions of the charge-specific percentile IM value for all targets and decoys. (d-e) Swarm plots of the number of peptides (d) and proteins (e) identified from all DDA PASEF runs. Figures a-c were generated using data from a single DDA PASEF run.

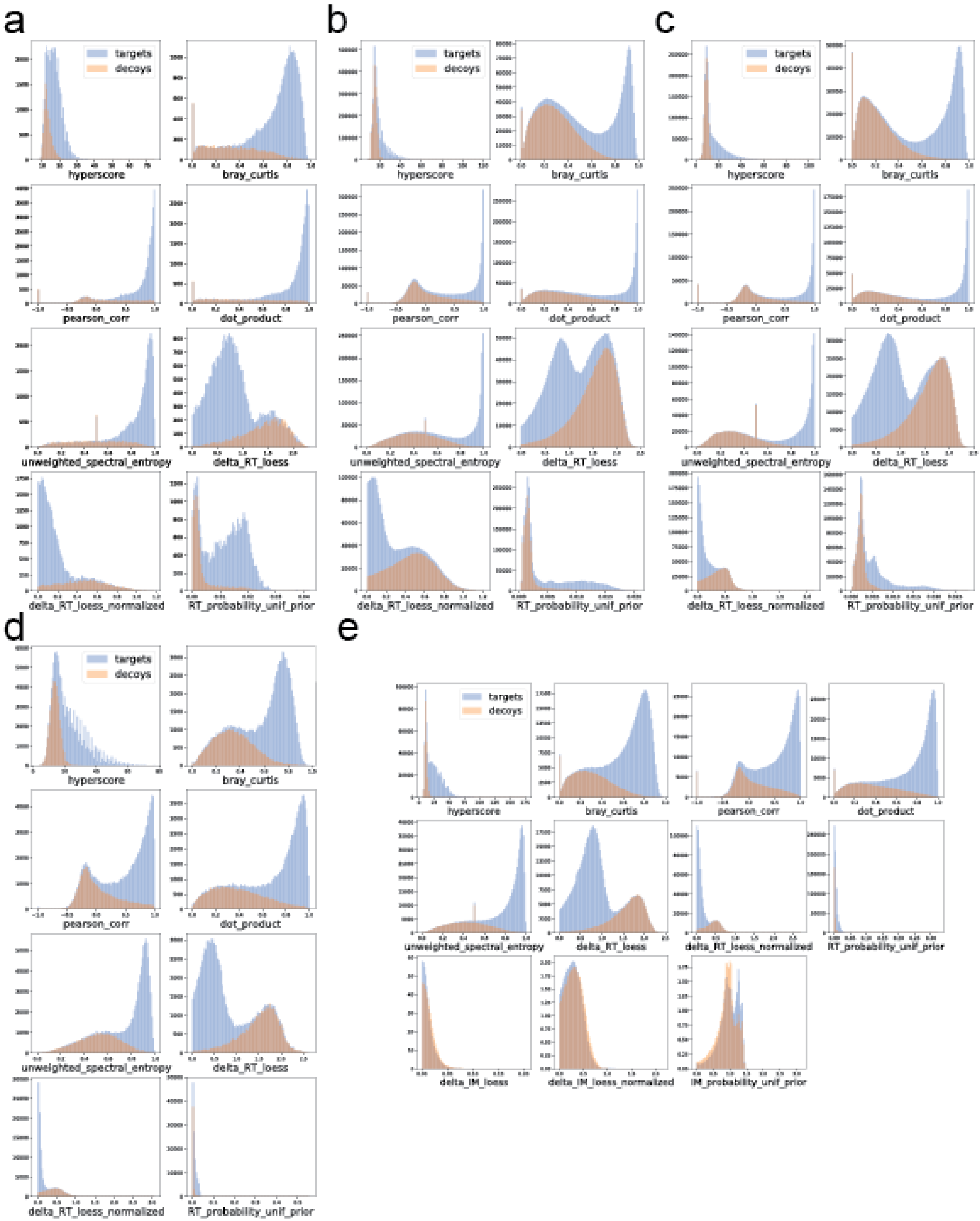


**Supplemental Figure 6.** PSM score distributions for MSBooster features. Target PSMs are in blue, decoy PSMs in orange. The datasets analyzed were HLA (a), melanoma processed by MSFragger DIA (b), melanoma processed by DIA-Umpire with MSFragger (c), single cell nanoPOTS (d), and timsTOF (e). RT and IM features were log-transformed to better visualize target-decoy separation using the conversion log10(value + 1). Spectral and RT figures have counts on the y-axis, IM has density.

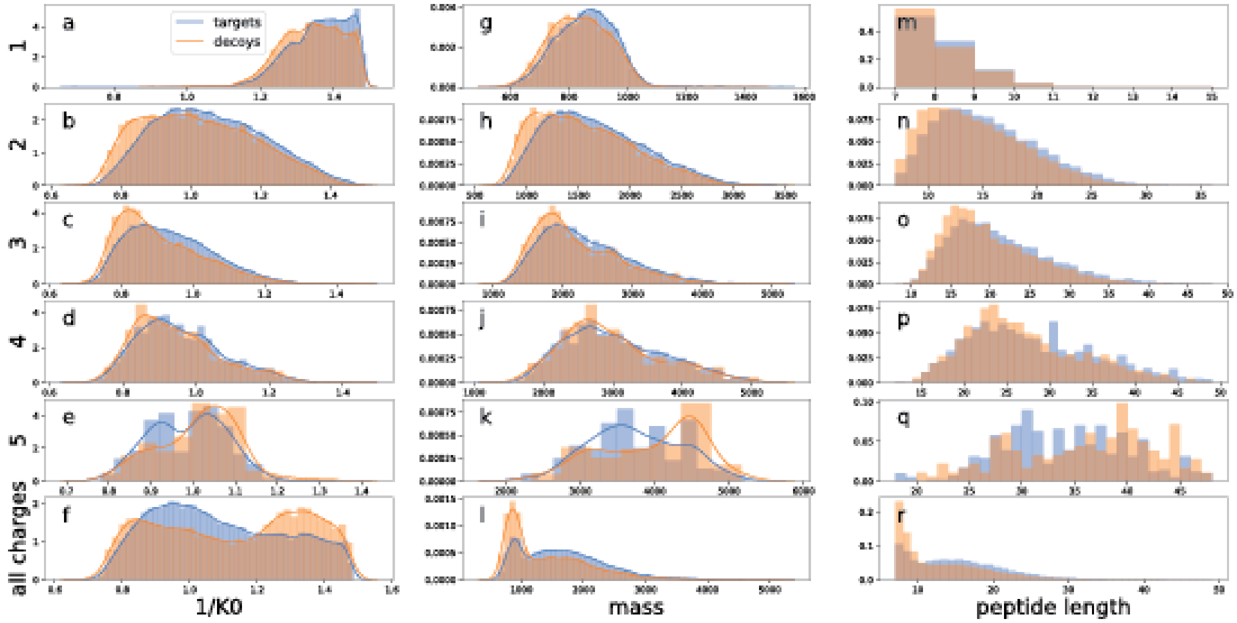


**Supplemental Figure 7.** Target-decoy PSM distributions for different charge states and features. All subfigures show relative density rather than PSM counts to better show target-decoy separation.

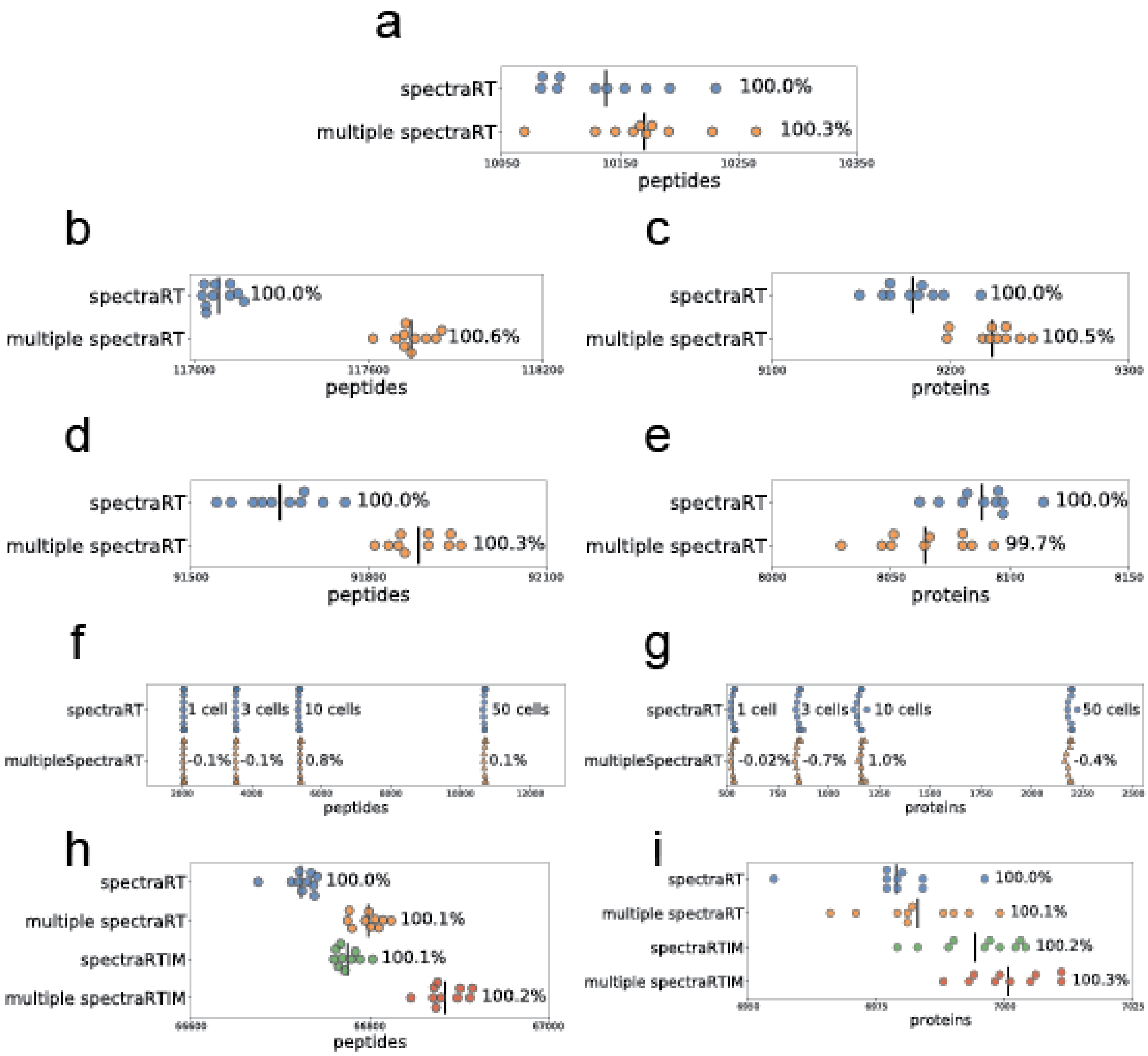


**Supplemental Figure 8.** Utility of multiple correlated features. Single features (unweighted spectral entropy + delta RT LOESS = spectraRT; unweighted spectral entropy + delta RT LOESS + IM probability uniform prior = spectraRTIM) are compared against multiple features (everything listed in Supplemental Note 1) for Percolator rescoring. Significance was calculated using independent t-tests with p<0.01. The numbers reported are for HLA peptides (a), melanoma peptides (b) and proteins (c) with MSFragger-DIA, melanoma peptides (d) and proteins (e) with DIA-Umpire and MSFragger, single cell peptides (f) and proteins (g), and timsTOF peptides (h) and proteins (i).

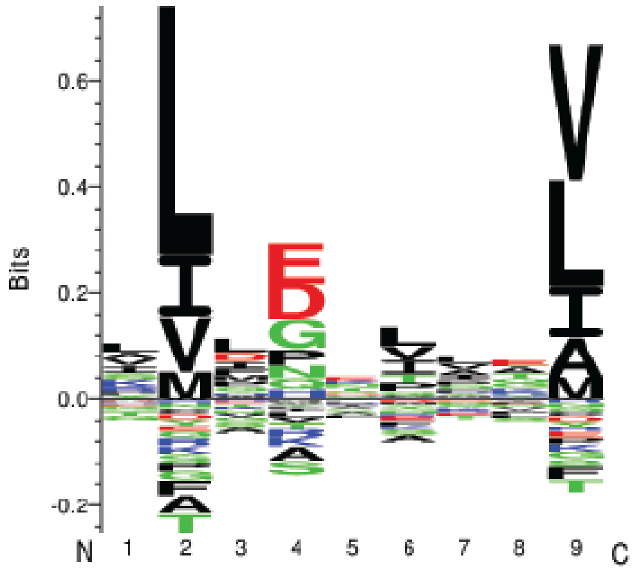


**Supplemental Figure 9.** HLA motif for new peptides from rescoring the HLA dataset with multiple correlated features.

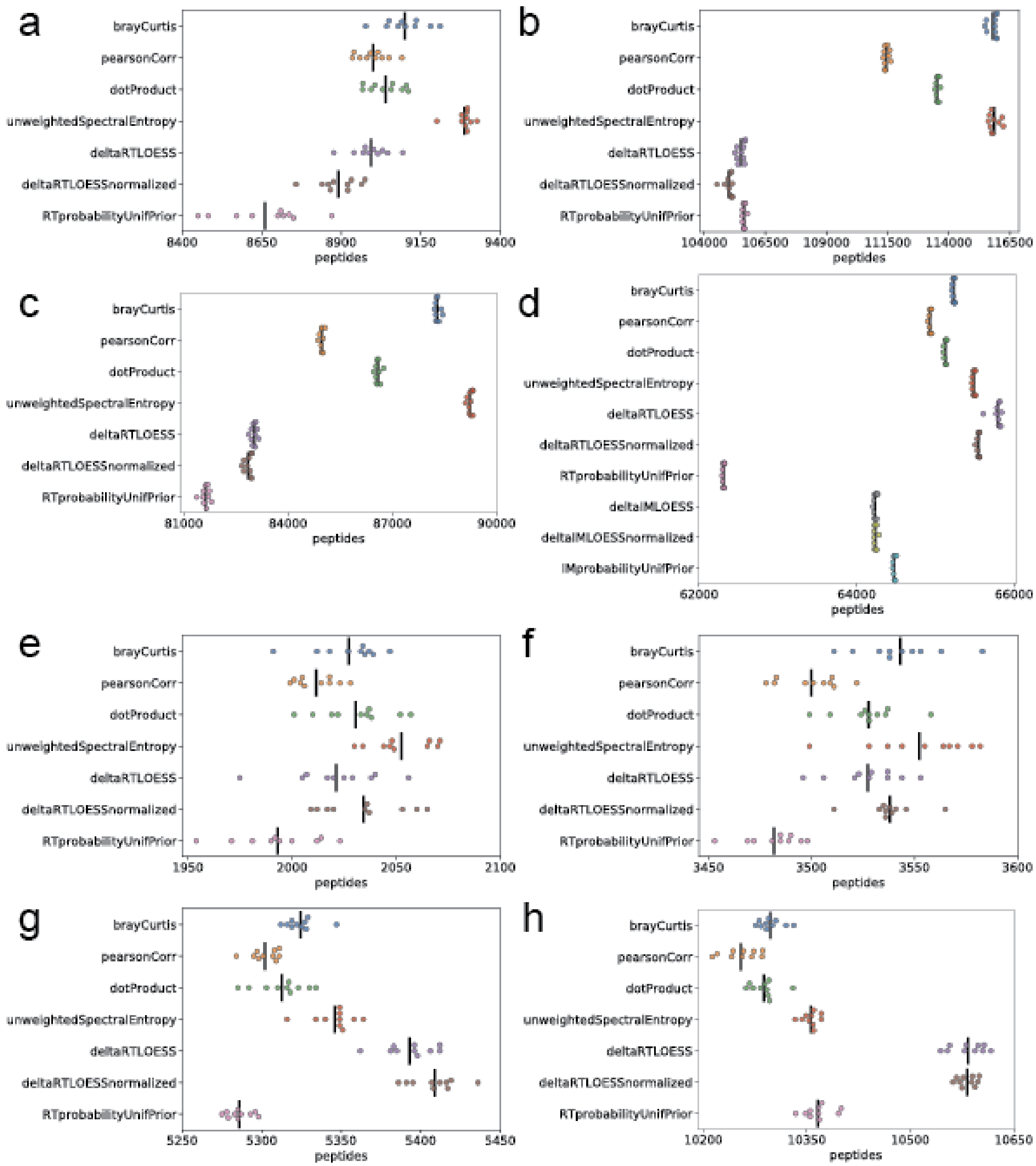


**Supplemental Figure 10.** Peptides reported when using different MSBooster features. Swarmplots for HLA (a), melanoma msfragger DIA (b), melanoma DIAU (c), timsTOF (d), 1 cell (e), 3 cells (f), 10 cells (g), 50 cells (h). Each dataset was processed 10 times by Percolator, each time with a different random seed 1-10. The black line indicates mean peptides reported over the 10 runs.
